# G2P: Using machine learning to understand and predict genes causing rare neurological disorders

**DOI:** 10.1101/288845

**Authors:** Juan A. Botía, Sebastian Guelfi, David Zhang, Karishma D’Sa, Regina Reynolds, Daniel Onah, Ellen M. McDonagh, Antonio Rueda Martin, Arianna Tucci, Augusto Rendon, Henry Houlden, John Hardy, Mina Ryten

**Author notes:** On behalf of the Neurology Genomics England Clinical Interpretation Partnership (Henry Houlden).

## Abstract

To facilitate precision medicine and neuroscience research, we developed a machine-learning technique that scores the likelihood that a gene, when mutated, will cause a neurological phenotype. We analysed 1126 genes relating to 25 subtypes of Mendelian neurological disease defined by Genomics England (March 2017) together with 154 gene-specific features capturing genetic variation, gene structure and tissue-specific expression and co-expression. We randomly re-sampled genes with no known disease association to develop bootstrapped decision-tree models, which were integrated to generate a decision tree-based ensemble for each disease subtype. Genes generating larger numbers of distinct transcripts and with higher probability of having missense mutations in normal individuals were significantly more likely to cause neurological diseases. Using mouse-mutant phenotypic data we tested the accuracy of gene-phenotype predictions and found that for 88% of all disease subtypes there was a significant enrichment of relevant phenotypic abnormalities when predicted genes were mutated in mice and in many cases mutations produced specific and matching phenotypes. Furthermore, using only newly identified genes included in the Genomics England November 2017 release, we assessed our gene-phenotype predictions and showed an 8.3 fold enrichment relative to chance for correct predictions. Thus, we demonstrate both the explanatory and predictive power of machine-learning-based models in neurological disease.

## Introduction

The last 10 years has seen amazing progress in the identification of genes associated with Mendelian forms of neurological diseases^1^. Each individual gene discovery is valuable for the patients affected, for the disease-based researchers and for the wider neuroscience community. However, these gene discoveries are perhaps even more important when considered collectively. First and foremost the growth of rare disease genetics has made it clear that dysfunction of the central and peripheral nervous system is a frequent outcome of genetic disorders with approximately 50% of all rare diseases (3000-3500 conditions) presenting with some form of neurological abnormality^2^. Furthermore, it has become clear that neurological disorders exhibit remarkable genetic heterogeneity. There are over 30 genes, which when mutated give rise to hereditary spastic paraplegia^3^ and over 80 genes associated with Charcot Marie Tooth disease^4^. Finally, it has become apparent that variable expressivity and atypical presentations are not the exception, but the rule when considering neurogenetic disorders^2^.

These observations have led us to hypothesise that there is something “special” about genes associated with neurogenetic disorders. Or more formally stated, we hypothesise that genes with significant effects on the structure and function of the nervous system share common characteristics not present amongst genes which when mutated do not affect the nervous system. Clearly identifying these common characteristics (assuming they are present) would be useful for two main reasons. Firstly, understanding the core characteristics of genes that give rise to neurological disorders would potentially help us to better understand the pathophysiology of at least a subset of disorders. Secondly, the identification of these common characteristics would allow us to scan the genome for other genes with the same key properties, but which have not yet been associated with neurological disease simply because the patient with this particular neurogenetic condition has not had appropriate genetic testing or the significance of the pathogenic variant could not be recognised. Since these predicted gene-disease associations would efficiently encapsulate prior knowledge, they could be used to prioritise genetic variants for further investigation when more standard analyses have failed to yield a result, an increasing clinical challenge.

While the identification of the core characteristics of a neuro-disease gene may seem an impossible task, there is an, ever increasing, quantity of public omic data, which has the potential to address this question. Over the last 3 years genome-wide data sets have become ever larger, more accessible, more consistent and more comprehensive both in terms of the depth of information offered and the quantity of data utilised. Public data sets include information on rare variant frequency across all genes^5^, gene and isoform structure^6^, and tissue-specific gene expression^7^. Hidden within these rich data sets could be key information regarding the relationship between genes and their functional effects in health and disease.

Machine Learning (ML) is well suited to the task of identifying such hidden phenomena and has been used successfully to address similar classification problems, such as the effects of genetic variants on exon inclusion^8^ and the prediction of genes associated with autosomal dominant disorders^9^. In this paper, we aim to use ML-based approaches to generate models to predict new genes associated with rare neurogenetic disorders. We base our learning experience on highly curated gene panels developed for the diagnosis of a wide range of neurological disorders. In this way, we aim to not only obtain novel insights into the pathophysiological processes underlying some neurogenetic disorders, but also provide predictive information, which can be rapidly translated for the benefit of patients.

## Results

### A genetic map of human neurogenetic disorders

Given that one of the major motivations of this study was the use of gene-phenotype association prediction as a means of improving the diagnostic yield of whole exome/ genome sequencing for rare neurogenetic disorders, we used gene panels generated by Genomics England (https://panelapp.genomicsengland.co.uk; March 31^st^ 2017) for “Neurology and neurodevelopmental disorders”. These gene panels are highly curated with over 45 clinicians and scientists contributing to the curation process (**Supplementary Table 1**). Furthermore, they are managed collectively to ensure quality across panels and aim to generate conservative “diagnostic-grade” gene sets, chosen because variants within the genes are capable of causing/explaining the specific neurological phenotype. Only gene panels with more than 10 “Green” (diagnostic-grade) genes were used in our analysis with the exception of the panel for intellectual disability, which contained 735 high confidence genes, but encompasses a phenotypically diverse disease set. This resulted in the inclusion of 25 panels in addition to a panel including all neuro-related genes together (**Supplementary Table 2**). Collectively this equated to 1140 unique genes. We noted a high degree of gene overlap amongst panels, prompting us to generate a genetic map of the inter-relationships between neurogenetic disorders based on “gene-sharing”. Interestingly, this “genetic map” broadly reflected recognised disease categories as exemplified by the clustering of neuromuscular disorders (Figure 1a).

**Figure 1.**
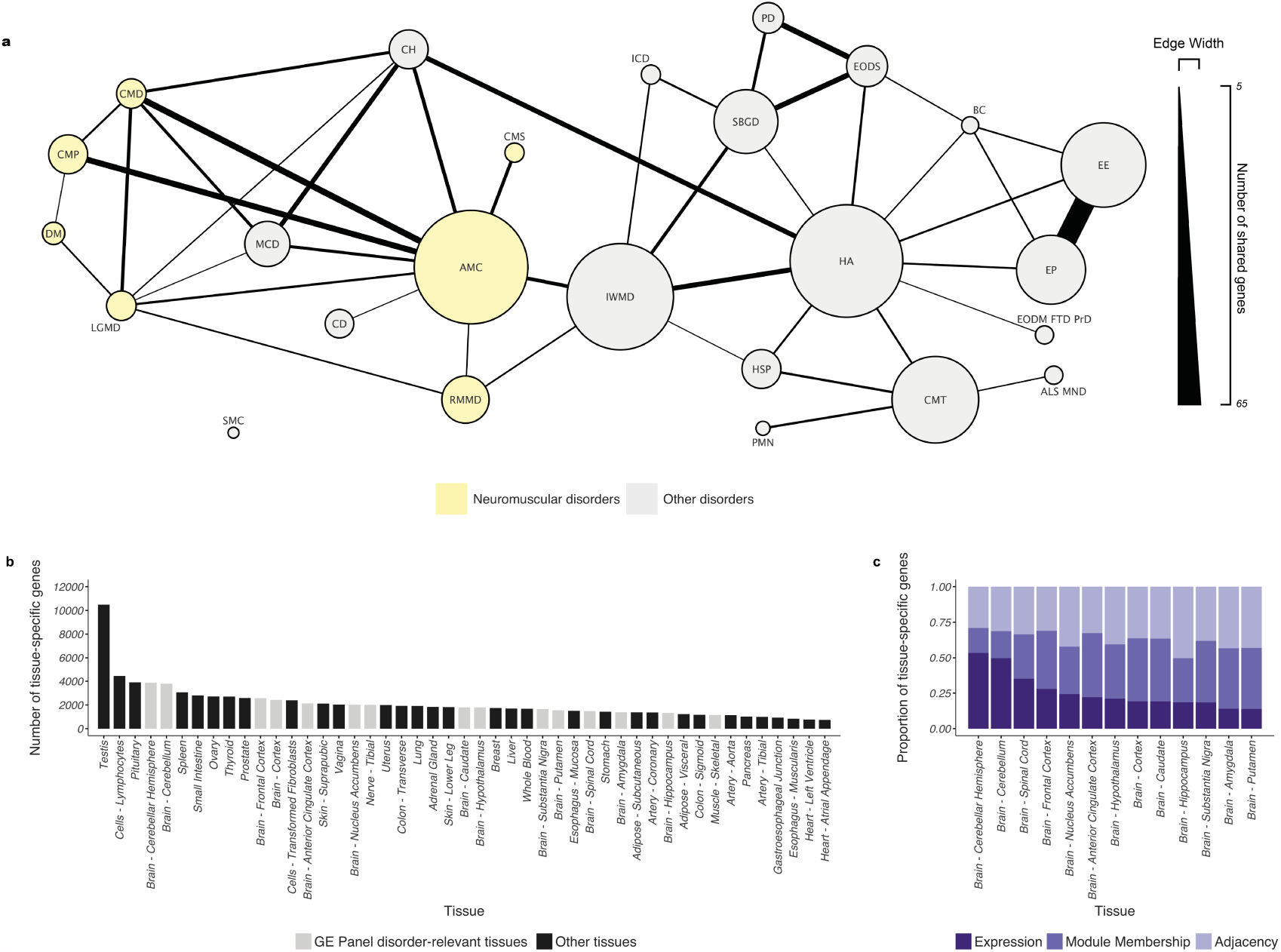
A genetic "map" of human neurogenetic disorders: **a**) The genetic overlap between Genomics England panels is displayed, such that each node represents 1 panel and a minimum of 5 genes is required for an edge. The radius of nodes and width of edges reflects the number of genes within each panel and the number of shared genes between panels respectively. **b**) Bar chart to show for each of the 47 GTEx tissues considered, how many genes expressed by the tissue show evidence of tissue-specificity on the basis of expression, adjacency or module membership values. We highlight in light grey tissues that appear as significant in the univariate test for any of these three features. **c**) Bar chart to show that brain tissues are characterised by high variability in the relative proportion of tissue-specific expression, adjacency and module membership. Cerebellar hemisphere shows the highest proportion of tissue-specific signals due to absolute expression, whereas putamen is characterised by the highest proportion of genes with tissue-specific co-expression. Genomics England panel names are shortened as follows: Amyotrophic lateral sclerosis motor neuron disease = ALS MND; Arthrogryposis = AMC; Brain channelopathy = BC; Cerebellar hypoplasia = CH; Cerebrovascular disorders = CD; Charcot Marie Tooth disease = CMT; Congenital muscular dystrophy = CMD; Congenital myaesthenia = CMS; Congenital myopathy = CMP; Distal myopathies = DM; Early onset dementia encompassing fronto temporal dementia and prion disease = EODM FTD PrD; Early onset dystonia = EODS; Epilepsy Plus = EP; Epileptic encephalopathy = EE; Hereditary ataxia = HA; Hereditary spastic paraplegia = HSP; Inherited white matter disorders = IWMD; Intracerebral calcification disorders = ICD; Limb girdle muscular dystrophy = LGMD; Malformations of cortical development = MCD; Paediatric motor neuronopathies = PMN; Parkinson Disease and Complex Parkinsonism = PD; Rhabdomyolysis and metabolic muscle disorders = RMMD; Skeletal Muscle Channelopathies = SMC; and Structural basal ganglia disorders = SBGD.

### Predictors derived from genomic and transcriptomic data are largely independent

Each of the 1140 known neurogenetic disease-related genes were considered for ML model development and were described together with all other protein coding genes (as defined by Ensembl version 72) using a set of “predictors”. We leveraged features extracted from large, public omic data sets to obtain these predictors of gene status (disease versus non-disease). We focused on genome-wide data sets, which do not incorporate any prior knowledge, either in the form of curation or biological/disease information, and which could provide gene-based metrics. This was a deliberate strategy in order to enable ML models to provide novel explanatory information not captured by current disease-related pathways or models.

More specifically, we collated the following three types of gene-specific information: i) gene-specific variant frequency, ii) gene and transcript structure, and iii) tissue-specific expression/co-expression of genes (**Online Methods & Supplementary Table 3**). Since deleterious variants are under negative selection, genes with less genetic variation than would be expected under a random mutation model might be supposed to be of particular disease-relevance. This information is already captured at the gene-level for specific mutation classes by the ExAC^5^ database parameters ExACpLI (the probability of a gene being intolerant of both heterozygous and homozygous loss-of-function variants), ExACpRec (the probability of being intolerant of only homozygous loss-of-function variants), and ExACpMiss (a Z-score reflecting the likelihood of being intolerant to missense mutations, see **Online Methods**). Conversely, tolerance to deleterious mutations as captured by ExACpNull (the probability of being tolerant of both heterozygous and homozygous loss-of-function variants) might be expected to be an attribute of non-disease genes. All four of the ExAC parameters were included as predictors in our learning set.

We also explored the possibility that the complexity of gene structure and transcription could be used to distinguish between disease and non-disease genes. We extracted or generated measures of structural complexity from GENCODE (Ensembl version 72) and the HEXEvent^10^ database. These measure of complexity included gene length, intronic length, the number of alternative transcripts generated, the number of possible unique exon-exon junctions, the number of alternative 3’ and 5’ sites used and the numbers of coding and non-coding genes overlapping with the gene of interest (**Online Methods**).

Given that disease-associated genes are often highly expressed in the tissues of relevance for the disease^11–16^, we included measures of tissue-specific expression in our learning data set. Using transcriptomic data generated by the GTEx^25^ project and covering 47 human tissues (including 13 brain regions, tibial nerve and skeletal muscle amongst others) we generated measures of tissue-specific gene-expression and co-expression (**Online Methods**). For each of the 47 human tissues we created a co-expression network using Weighted Gene Co-expression Network Analysis^17^ optimized by k-means^18^. This provided estimates of each gene’s global and local connectivity in relation to all other expressed genes in the tissue (as captured by the terms “adjacency” and “module membership”, **Online Methods**). This allowed us to generate measures of the tissue-specific network properties of each gene and created an additional 141 binary predictors.

Interestingly, inspection of this set of predictors demonstrated that the absolute number of genes with evidence of tissue-specific expression/co-expression was highly variable across tissues (Figure 1b). While 10,483 genes showed testes-specific gene expression or co-expression, only 1699 genes were expressed in a tissue-specific manner in liver. Furthermore, the relative importance of tissue-specific gene expression, as distinct from co-expression (Figure 1b & 1c) was also very variable across the 47 GTEx tissues. If we focus on brain tissue, whereas 49.7% of tissue-specific genes for cerebellum were defined as cerebellar-specific due to high absolute expression in the tissue, in putamen the majority of specific gene expression was detected through the unique local and global network properties of genes (86.12%, Figure 1c).

Finally, we explored all 154 predictors across the major classes, namely genetic, structural and expression-based predictors, for evidence of correlation amongst predictors (**Supplementary Table 4**). Predictors within a major class (genetic variation, gene structure, gene expression, gene co-expression) were more correlated to each other than to predictors of a different class suggesting that the major predictor classes used were capturing orthogonal types of gene-specific information (Figure 2).

**Figure 2.**
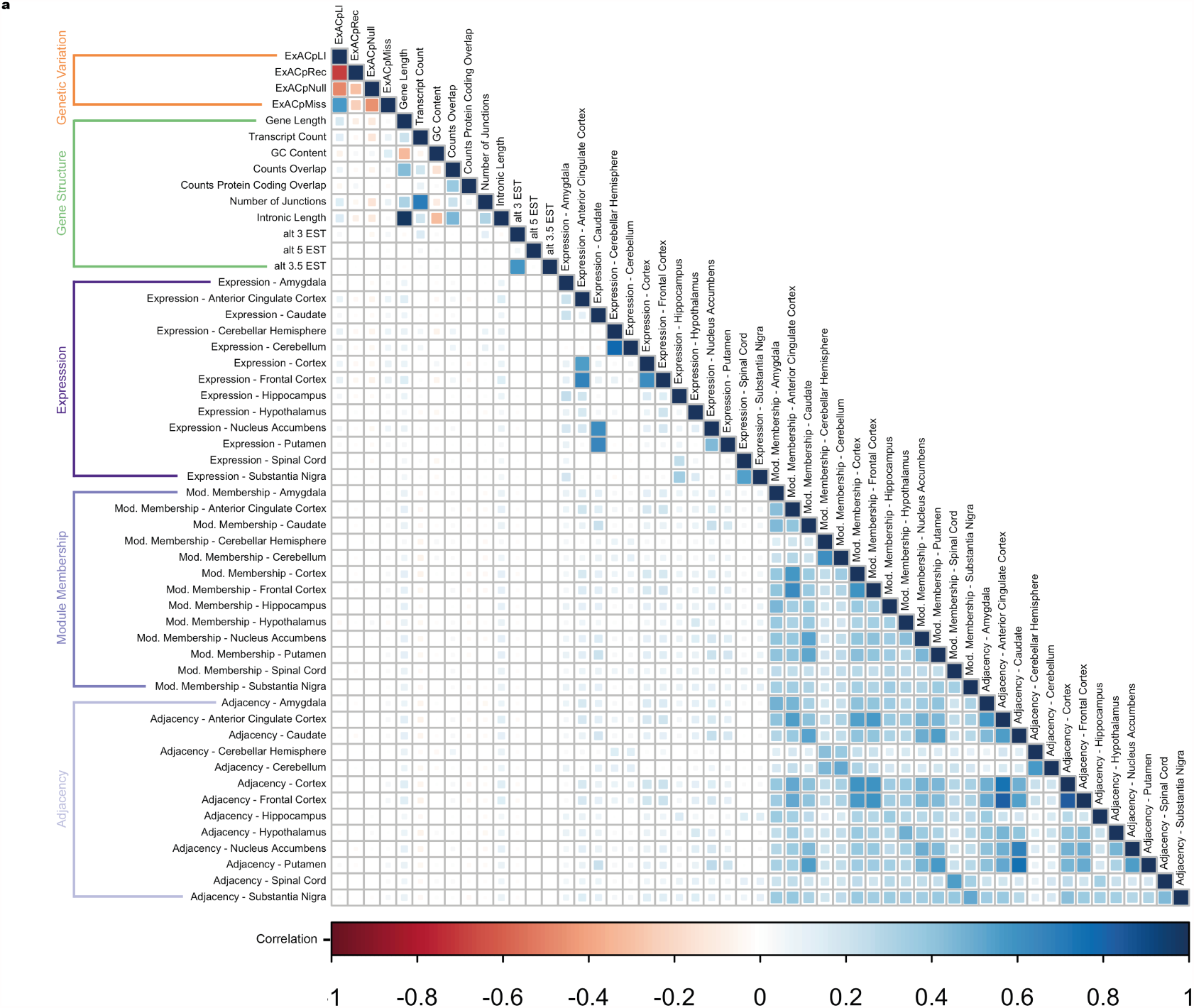
The three major types of predictors, namely genetic variation, gene and transcript structure, and gene expression/co-expression, are largely independent: Correlation matrix plot to show the Pearson’s correlation coefficients between the predictor categories, namely genetic variation (orange), gene and transcript structure (green), tissue-specific gene expression (dark blue), tissue-specific module membership (blue) and tissue-specific adjacency (light blue).

### Measures of gene complexity and intolerance to missense variants distinguish between disease and non-disease genes across most disorders

As described above we generated 154 potentially useful predictors of disease-gene state for each gene within the human genome. In the first instance we wanted to find out whether there were any statistical differences in predictor values between established disease-related and non-disease genes using single predictors alone. This testing was performed for each of the 25 disease gene panels separately and for all predictors whether numerical or categorical in nature (**Online Methods**).

Using this approach we found significant differences (FDR corrected p-value <0.05) in at least a single predictor for 22 of the 25 disease gene panels (88.0%, **Supplementary Table 5**). Of the three main types of gene-specific predictors analysed (gene-specific variant frequency, gene and transcript structure, and tissue-specific expression/co-expression of genes), we found that gene and transcript structure appeared to be the most useful. Of all 25 disease gene panels 72.0% showed a significant difference in predictor values for at least one of the measures of gene and transcript structure. Considering transcript count alone (the number of transcripts produced by a gene), we found that 56% of all the disease panels (minimum FDR corrected p-value = 5.22 × 10^−18^ for “Epilepsy-Plus”) had a significantly higher transcript count for disease-related as compared to non-disease genes (Figure 3a, **Supplementary Table 5**). The second most important category of predictor was gene-specific variant frequency with 64.0% of all disease panels showing a significant difference in predictor values for at least one measure, with ExACpMiss accounting for the largest proportion of panels (40.0%, Figure 3b).

**Figure 3.**
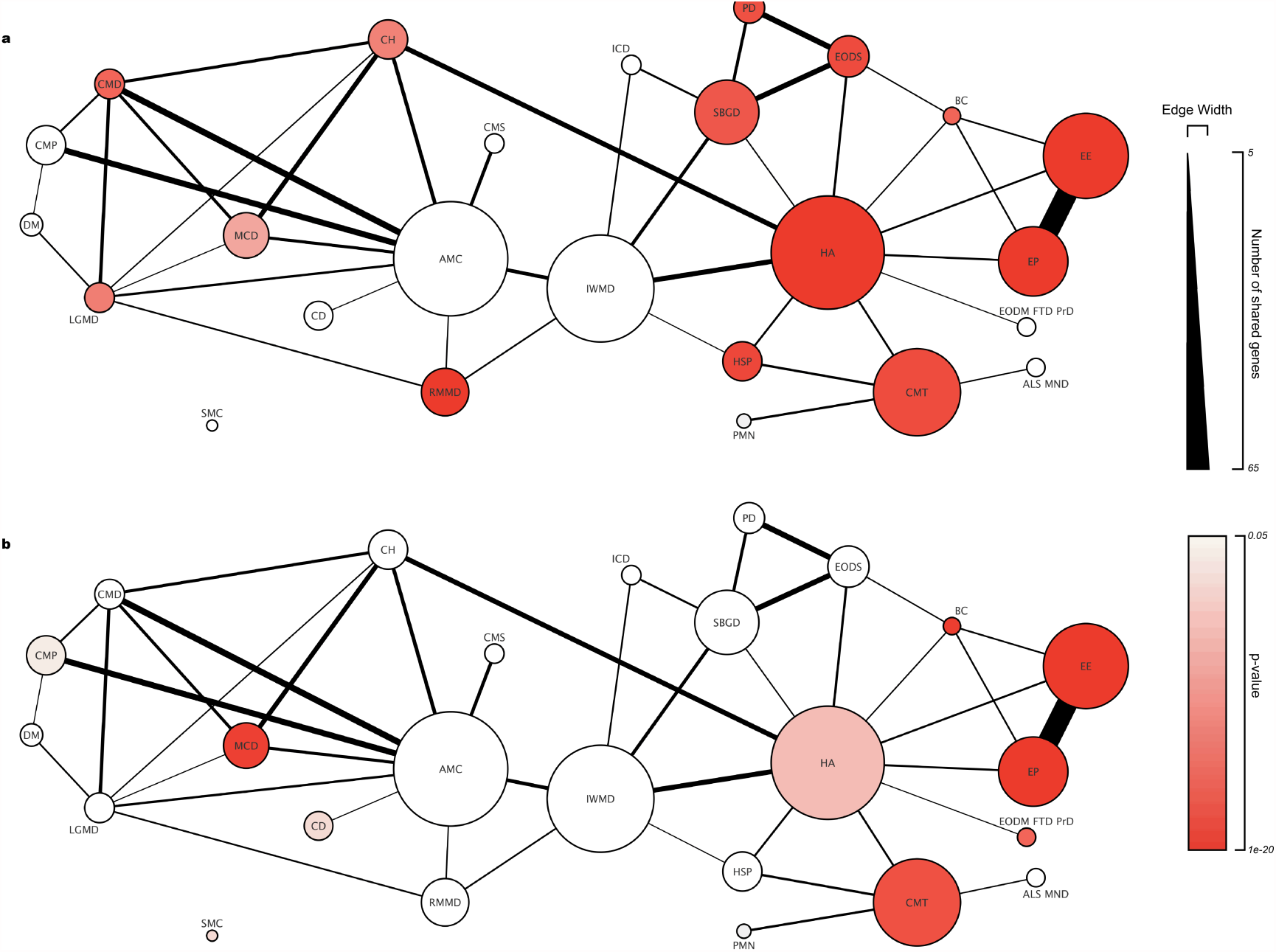
Transcript complexity and reduced tolerance to missense mutations are a distinguishing feature of genes associated with many types of rare neurogenetic disorders: **a**) Plot showing that of the 25 Genomics England panels considered 56% had a significant p-value (red scale) on comparing transcript count values between genes in the panels and other protein coding genes. **b**) Plot showing that of the 25 Genomics England gene panels considered 40% had a significant p-value (red scale) on comparing ExACpMiss values between genes in the panels and other protein coding genes. Genomics England panel names are shortened as follows: Amyotrophic lateral sclerosis motor neuron disease = ALS MND; Arthrogryposis = AMC; Brain channelopathy = BC; Cerebellar hypoplasia = CH; Cerebrovascular disorders = CD; Charcot Marie Tooth disease = CMT; Congenital muscular dystrophy = CMD; Congenital myaesthenia = CMS; Congenital myopathy = CMP; Distal myopathies = DM; Early onset dementia encompassing fronto temporal dementia and prion disease = EODM FTD PrD; Early onset dystonia = EODS; Epilepsy Plus = EP; Epileptic encephalopathy = EE; Hereditary ataxia = HA; Hereditary spastic paraplegia = HSP; Inherited white matter disorders = IWMD; Intracerebral calcification disorders = ICD; Limb girdle muscular dystrophy = LGMD; Malformations of cortical development = MCD; Paediatric motor neuronopathies = PMN; Parkinson Disease and Complex Parkinsonism = PD; Rhabdomyolysis and metabolic muscle disorders = RMMD; Skeletal Muscle Channelopathies = SMC; and Structural basal ganglia disorders = SBGD.

While tissue-specific expression/co-expression of genes could be highly discriminatory predictors, this was restricted to small sets of related disorders. For example, specific expression in skeletal muscle was a highly significant predictor for a range of primary muscle disorders, including “Distal myopathies” (p-value 2.78 × 10^−32^), “Congenital myopathy” (p-value 2.35 × 10^−22^) and “Rhabdomyolosis and metabolic muscle disease” (p-value 2.94 × 10^−15^). Similarly, specific expression in the frontal cortex could be used to distinguish between epilepsy-related and non-disease genes (p-value 1.26 × 10^−10^ and p-value 3.53 × 10^−7^ respectively).

Thus, this analysis demonstrated that while genes associated with neurological diseases have common characteristics of broad relevance with complexity of transcript structure being the single most important discriminatory genic feature, there are also features operating in a more disease-specific manner.

### ML-based classifiers for disease gene prediction can be optimized for accuracy at the expense of genome coverage

The major aim of this study was to generate machine learning classifiers, based on genetic, structural and expression-based predictors, to distinguish between genes which when mutated produce a specific neurological phenotype versus those that do not. We wanted to create such classifiers for each of the 25 disorders as defined by Genomics England. This was challenging because not only do we require classifiers capable of providing explanatory as well as predictive information, but the generation process had to be robust despite the large disparity in the size of disease (mean number of genes per panel = 45, range 18-111) and non-disease gene sets (4913 genes, 91.59% of the total set of learning examples).

Our strategy was to address these issues in two separate steps. In the first step we evaluated a range of learning paradigms to select one with a reasonably good trade-off between predictive and explanatory power. In the second step, we created an ensemble composed of many single ML models based on the learning paradigm of choice (as identified in step 1). More specifically, the first step involved testing a range of ML approaches covering the main supervised learning paradigms and available through the Caret R package^19^ (**Online Methods**). We obtained ROC estimates (a single measure, which conveniently integrates sensitivity and specificity) for each algorithm applied to each of the 25 gene panels (**Supplementary Figures 1 & 2**). On the basis of these results, we selected J48^20^ (**Online Methods**), a decision tree-based algorithm, due to its explanatory capabilities and high predictive power.

In the second step, we addressed the imbalance of positive and negative learning examples (average ratio of positive to negative learning examples = 1:109) by creating an ensemble of J48 models obtained from learning datasets with equal numbers of positive and negative gene examples. We created each learning dataset by randomly re-sampling non-disease genes (Figure 4a), while keeping all disease genes unchanged. In this way for every disease category we generated 200 separate J48 trees, each trained using the same disease gene set, but a different, randomly generated “non-disease” gene set. We integrated the 200 classifiers generated per panel using a voting approach and operated a simple majority to determine the outcome for a given gene. This meant that for a given gene, we predicted a gene-phenotype relationship as present if most of the classifiers “voted disease” and absent if most of the classifiers “voted non-disease”. We assessed whether the integration of disease classifiers to generate a classifier ensemble improved performance. We found that for all 25 disease categories using a classifier ensemble improved the ROC values. On average there was a 4.1% improvement in ROC values (range of % improvement = 0.6% - 7.6%), relative to using a single J48 tree (**Supplementary Figure 3**).

**Figure 4.**
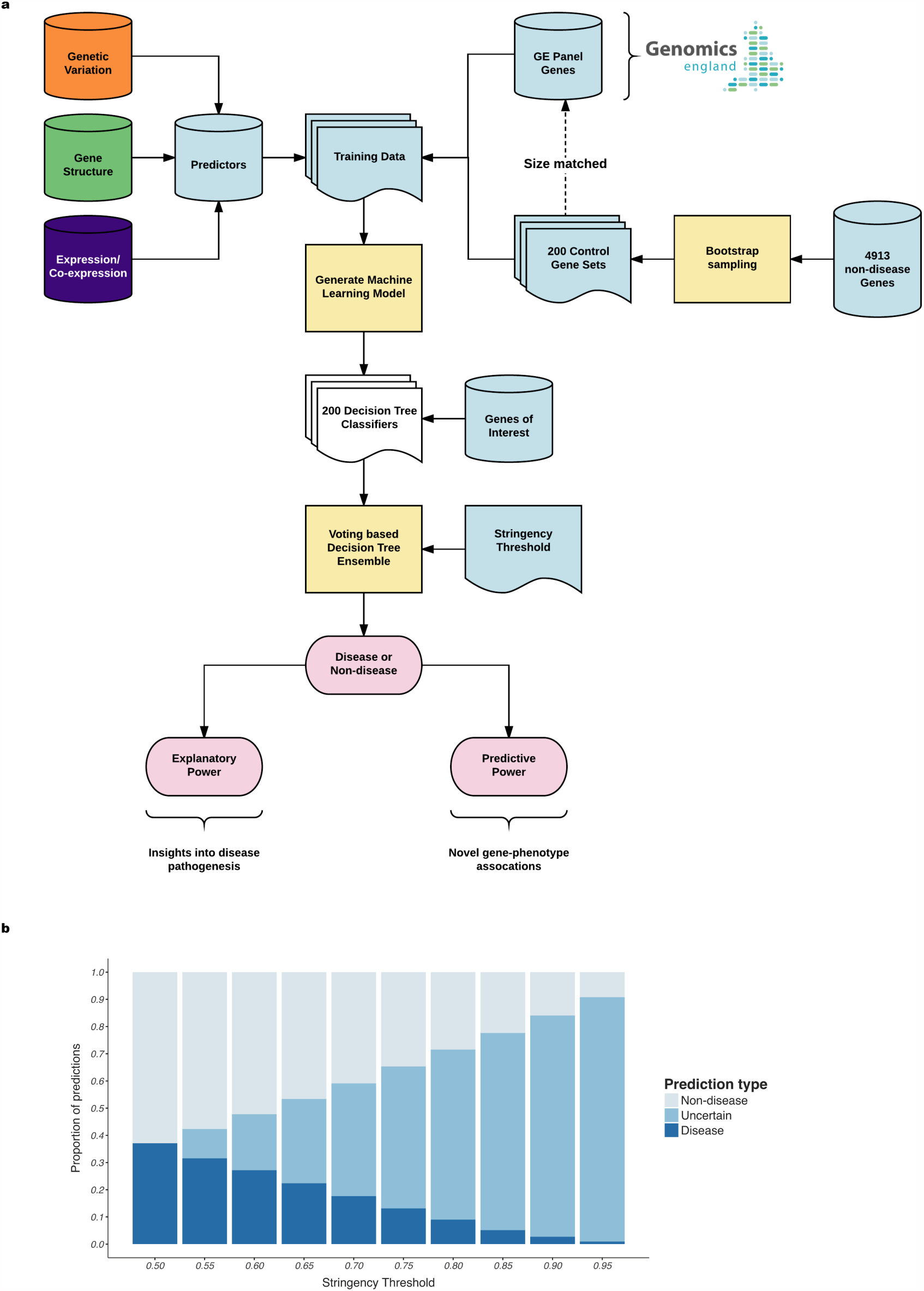
Construction and calibration of ML classifier ensembles for gene-phenotype prediction: **a**) Diagram to show the workflow used to generate ML classifier ensembles. These ensembles are based on the creation of 200 learning data sets per a disease panel. Each data set is generated by resampling the control gene set to maintain a 1:1 ratio of disease and control genes. The final model of 200 decisition trees are integrated into a voting scheme that can be used to prediction novel gene-phenotype associations with an estimate of confidence (bottom right part) and to identify the genic features contributing most commonly to the prediction process (bottom left part of the figure). **b**) Plot to show the effect of applying a stringency parameter on ML predictions generated from decision tree ensembles. As stringency values increase (x-axis) the relative proportions of "Disease", "Non-disease" and "Uncertain" also change. In particular, the proportion of uncertain predictions increases.

Furthermore, the integration of disease classifiers using a simple voting approach allowed us to adjust the stringency of our classification system by changing the percentage of votes required (termed the “stringency parameter, s”) to confidently assign a gene as “disease” or “non-disease”. In this way, we converted a 2-output ensemble into a 3-output ensemble adding “uncertain” as the third possible outcome, used when there was not enough agreement amongst votes. To illustrate the effect of the stringency parameter (“s”) on prediction of “disease” genes, we tested a range of “s” values across all disease ensembles and all protein coding genes (Figure 4b). This demonstrated that as the stringency increased, the proportion of genes that could be classified either as “Disease” or “Non-disease” decreased. Thus, increasing our confidence in gene-phenotype predictions, comes at the cost of genome coverage with the proportion of genes for which we generate an “uncertain” outcome rising rapidly. Therefore, depending on the downstream use of classifiers (e.g. for use in a diagnostic versus a research setting) we would envisage situations where it would be sensible to adjust the stringency. For this reason we have released a web interface where predictions can be obtained at variable levels of stringency for all diseases (www.XXXXXX).

### Predicted genes are enriched for relevant gene ontology terms and cell-specific markers

After removing genes used to train classifiers, we applied all 26 of the classifier ensembles to the entire set of human protein coding genes as defined within GENCODE (16,012). Using a stringency of 0.9 for each classifier (i.e. predict Disease or Non-disease only when the majority of identical predictions is equal or above 90%), we generated 4,834 new gene-phenotype predictions (Figure 5a, **Supplementary Table 6**). On average, each classifier predicted 272 new disease genes (equating to 1.7% of the gene classifications attempted). As would be expected from the high numbers of overlapping genes across disease panels, there was a high degree of sharing amongst gene predictions from different classifier ensembles. Nonetheless, all classifier ensembles generated predictions which were disease-specific (Figure 5a). We noted that the number of new “disease” predictions was positively correlated (Pearson correlation = 0.71, p-value < 4.03 × 10^−5^) with the size of the original disease panel (i.e. the number of genes already known to be associated with disease, Figure 5a). No significant correlation was detected when we considered non-disease predictions and this is consistent with our expectation that the success of ML-based classifiers depends on the adequacy of positive training examples.

**Figure 5.**
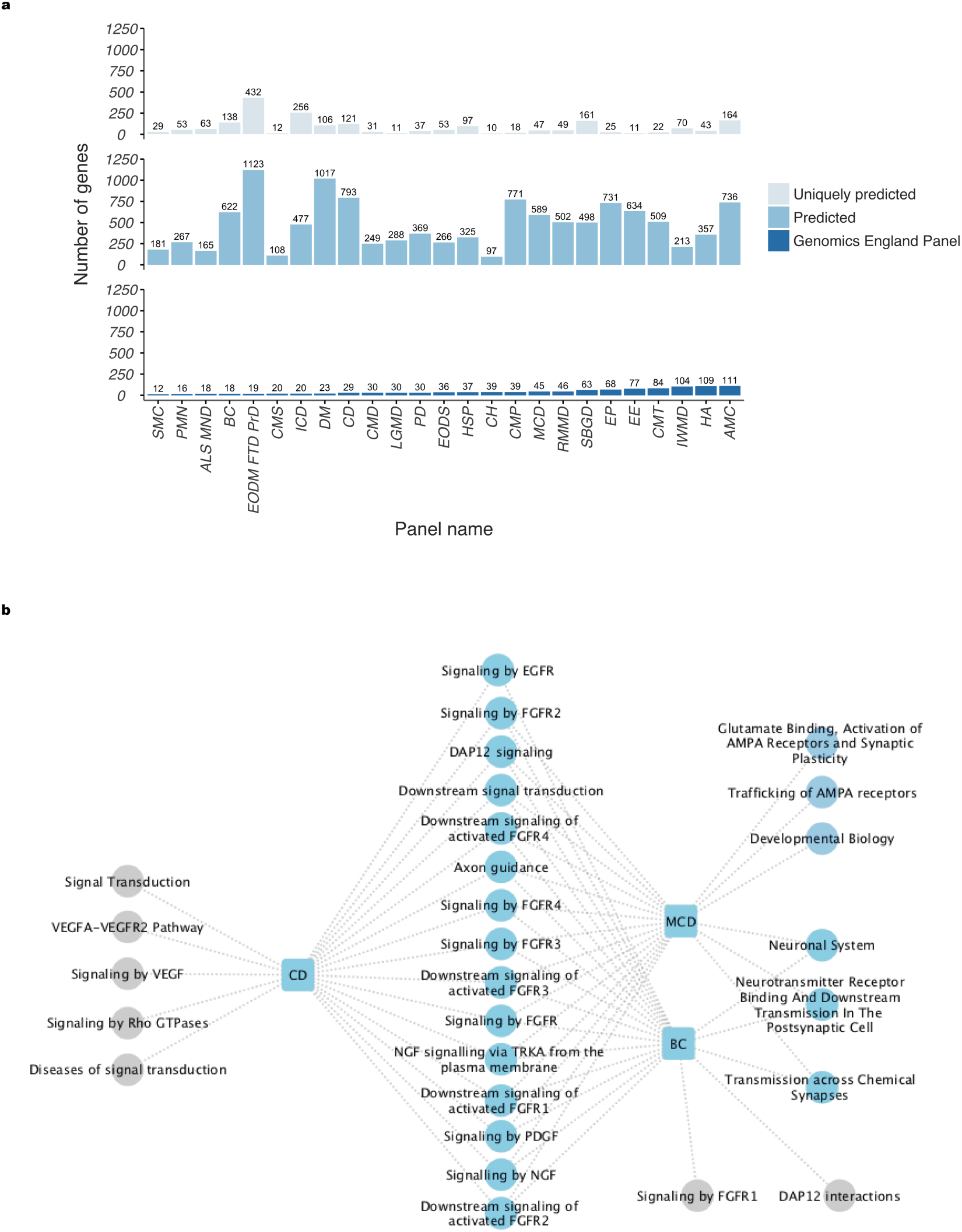
Properties of gene-phenotype predictions generated by disease-specific ML classifier ensembles: **a**) The number of number of genes contained within each Genomics England disease panel and used for the generation of ML classifier ensembles is shown in the lower panel (positive example set). The total number of new genes predicted per panel using a stringency of 0.9 is shown in the middle panel. The upper panel shows the number of predicted genes per panel, which were exclusively predicted by the disease-specific classifier. **b**) Plot to show that genes predicted to cause malformations of cortical development are significantly enriched for relevant biological processes. REACTOME pathways (circles) with the most significant enrichments amongst genes predicted to cause "Malformations of cortical development" (labelled rounded square) are shown. REACTOME pathways which are also highlighted by testing for enrichment of predicted genes for other disorders, namely " Cerebrovascular disorders" and "Brain channelopathies”, are also displayed (shared enrichments = blue circle; enrichments unique to either “Cerebrovascular disorders" or "Brain channelopathies" = grey circles).

Given that no cellular, pathway or disease-based information was used within the learning data, we wanted to investigate the biological characteristics of our predicted genes. We did this first by assessing predicted genes for enrichment of gene ontology terms^21^, as well as REACTOME^22^ and KEGG^23^ pathway annotation (Figure 5b, **Online Methods**). Using this approach and considering disease gene predictions produced by each ensemble separately, we identified 2,345 unique significant terms (at an FDR corrected p-value of <0.05; **Supplementary Table 7**). Interestingly gene predictions made by the “all in one” classifier (based on the entire set of disease genes) was highly enriched for terms relevant to the nervous system with the top REACTOME pathway terms being “Neuronal System” (FDR-corrected p-value = 5.12 × 10^−15^), “Transmission across Chemical Synapses” (FDR-corrected p-value = 5.12 × 10^−9^) and “Axon guidance” (FDR-corrected p-value = 3.57 × 10^−8^, **Supplementary Table 7**).

Inspection of disease-specific enrichments also demonstrated the relevance of predicted genes. For example, amongst genes predicted by the classifier for “Malformations of cortical development” there were significant enrichments for 966 terms alone. Focusing on REACTOME pathway enrichments, we identified a large number of significant terms relating to growth factor signalling and neuronal migration (Figure 5b, **Online Methods**). In the case of the former, this included “Downstream signaling of activated FGFR2” (FDR-corrected p-value = 2.21 × 10^−9^), a signalling system which has been specifically implicated in the function of outer radial glia^24^. This analysis also highlighted genes involved in axon guidance (FDR corrected p-value = 3.93 × 10-16) and synapse formation (“Transmission across Chemical Synapses”, FDR-corrected p-value = 2.49 × 10^−16^; “Glutamate Binding, Activation of AMPA Receptors and Synaptic Plasticity”, FDR-corrected p-value = 4.77 × 10^−12^), consistent with the known importance of neuronal migration abnormalities in the pathophysiology of cortical malformations^25^. Furthermore, a closer investigation of the sharing of significant REACTOME terms across predicted gene sets (generated with different classifiers) revealed the possibility of shared pathophysiological processes underlying disorders. For example, enrichment analyses for genes predicted by both the “Cerebrovascular disease” and “Malformations of cortical development” classifiers, were highly enriched for genes involved in growth factor signalling (Figure 5b), despite there being no overlap in the genes used as positive learning examples for classifier generation.

We also examined predicted gene sets for evidence of enrichment of cell-specific gene markers. This analysis was performed using gene expression signatures derived from RNA sequencing of purified cell types isolated from mouse cerebral cortex^26^ and covered astrocytes, neurons, oligodendrocyte precursors, newly formed oligodendrocytes, myelinating oligodendrocytes and endothelial cells. Using this approach and considering disease gene predictions produced by each ensemble separately, we identified 22 significant cell-specific enrichments (FDR < 0.05, **Supplementary Table 8, Online Methods**). In many cases, these enrichments were consistent with the phenotypic features of the disorder. For example, genes predicted by the classifier for “Cerebrovascular disorders” were significantly enriched for endothelial cell markers, while gene predictions for “Inherited white matter disorders” were enriched for markers of oligodendrocyte precursor cells.

Thus, these analyses provide evidence that the classifiers we generated capture some important elements of the biology of neurogenetic disorders and so are capable of generating predicted genes, which share those key biological properties.

### Disrupting predicted genes tends to produce neurologically-relevant phenotypes in mice

In order to explore the accuracy of gene predictions further, we made use of the MGI Mouse-Phenotype Database^27^ to determine whether disruption of mouse orthologues of predicted human genes produced phenotypes relevant to the neurological diseases studied. Considering the genes predicted by each of the 26 classifiers separately, we identified 1588 unique significantly enriched Mammalian Phenotype Ontology (MPO) terms of which 28.7% were directly relevant to the central and/or the peripheral nervous system (defined as sub-terms of MP:0005386, behaviour/neurological phenotype; MP:0003631, nervous system; MP:0005369, muscle phenotype). Using genes predicted by the “all in one” classifier (based on the entire set of disease genes), we identified 173 terms with significant enrichment in mouse model systems (**Supplementary Table 9**), with the most significant MPO terms being “abnormal synaptic transmission” (FDR-corrected p-value = 1.20 × 10^−10^) and “abnormal nervous system physiology” (FDR-corrected p-value = 1.22 × 10^−8^).

We also explored the enrichment of MPO terms amongst mouse models relevant to disease-specific classifiers. Despite the challenges inherent in this analysis, namely that this is a cross-species analysis using data from mice with both spontaneous and targeted mutations of multiple types, we were able to identify examples where disruption of predicted genes mirrored the expected disease phenotype with precision. For example, using the Genomics England Panel for “Malformations of cortical development”, which consists of 45 genes, we predicted a further 589 genes (using stringency 0.9). Of the 585 relevant mouse orthologues, mouse phenotypic data was available for 427 genes. Using this information we found that disruption of these genes in mouse models resulted in phenotypes highly enriched for “abnormal cerebral hemisphere morphology” (FDR-corrected p-value = 1.14 × 10^−13^), and more specifically “abnormal cerebral cortex morphology” (FDR-corrected p-value = 2.72 × 10^−7^).

### Genes predicted using ML-based classifiers are enriched for true gene-phenotype associations in humans

Clearly the most important means of assessing the value of ML-based classifiers for the prediction of genes contributing to neurogenetic conditions, is the identification of mutations in predicted genes in patients with matching neurological phenotypes. We assessed this systematically by making use of the regular re-versioning of Genomics England panels and the continuous updates available through the Online Mendelian Inheritance in Man catalogue (OMIM, www.OMIM.org). In the case of the former, we identified all high confidence gene-phenotype relationships added to PanelApp after the March 31^st^ 2017 version release and until November 1^st^ 2017. In the case of the latter, we collated all gene-phenotype associations included between 1^st^ April 2017 and 5^th^ January 2018. After excluding all OMIM gene-phenotype associations classed as provisional, we manually curated the remaining associations to ensure that the phenotypes could be mapped accurately to a Genomics England disease panel. Using this approach we identified a total of 32 new, high confidence gene-phenotype associations for which the relevant ML-based classifier provided a “Disease” or “Non-disease” prediction (applying a stringency of 0.9). We correctly predicted “Disease” status in 17 cases with the remaining 15 genes predicted as “Non-disease” (**Supplementary Table 10**, ratio of “Disease”/”Non-disease” = 113.3%). Given that on average the ratio of “Disease” to “Non-disease” predictions amongst the relevant ML-based classifiers obtained on the whole protein coding genome is 13.9%, this represented an 8.2 fold enrichment over chance.

Furthermore, exploring individual examples enabled us to recognise the explanatory value of classifier ensembles. For example, using the ML-based ensemble for “Cerebellar hypoplasia”, which was trained with 39 known disease genes, we correctly predict that mutations in *TBC1D23* are a cause of the disorder^28^. Interestingly, inspection of all the decision trees demonstrated that the two most important predictors were cerebellum-specific gene expression and gene complexity, as captured by the number of transcripts produced by a gene (Figure 6a, **Online Methods**). In fact, accounting for both the usage of a predictor (Appearance Index) and its depth (a measure of its average proximity to the root) across all 200 decision trees, demonstrated that these were more important predictors than gene-specific measures of variant frequency, such as ExACpLi and ExACpMiss scores, which are more commonly used to assess the pathogenicity of variants. Consistent with these findings we noted that *TBC1D23* has higher mRNA expression in cerebellum as compared to other brain regions not only in the adult brain but throughout most of brain development (Figure 6b). Furthermore, it produces a higher than expected number of transcript count when considering all protein coding genes (Figure 6c).

**Figure 6.**
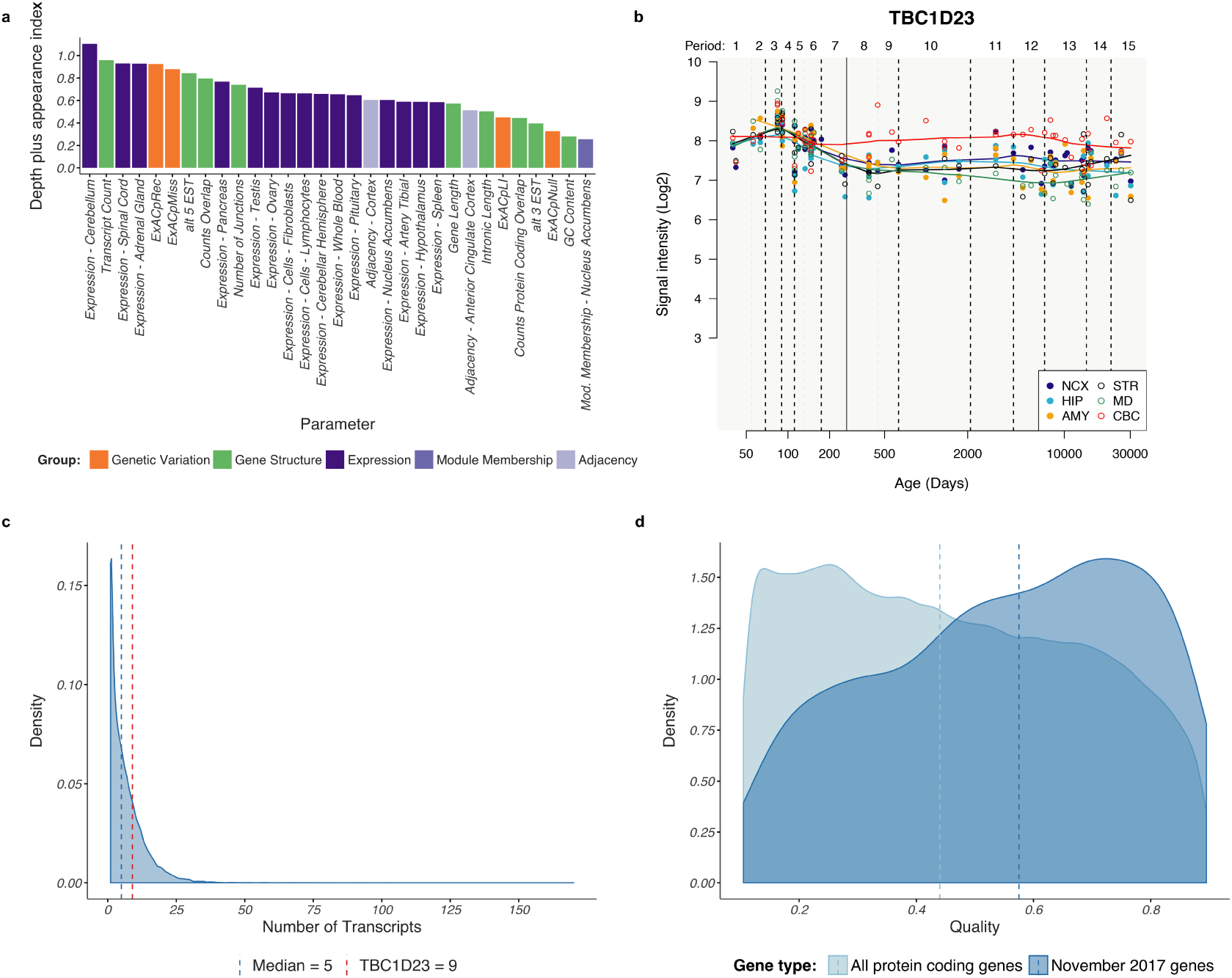
Evidence for enrichment of true gene-phenotype associations amongst gene predictions made by disease-specific classifiers: **a**) Cerebellum-specific gene expression and transcript complexity are the most important predictors used within the “Cerebellar hypoplasia” classifier ensemble. On the x-axis we plot each of the predictors used. Predictor importance is plotted on the y-axis, as defined as the sum of the appearance index of a predictor (the percentage of times it is used to predict “Disease” across the 200 decision trees which comprise the ensemble classifier, range = 0.0-1.0) and the depth of a predictor (a measure of its average proximity to the root, range = 0.0 – 1.0). This plot demonstrates that the most important predictors in the “Cerebellar Hypoplasia” ensemble classifier were cerebellar-specific gene expression and transcript count (the number of transcripts produced by a gene as annotated in GENCODE version 72). **b**) Graph to show mRNA expression levels for *TBC1D23*^29,30^ in 6 brain regions during the course of human brain development, based on exon array experiments and plotted on a log2 scale (y axis). The brain regions analyzed are the striatum (STR), amygdala (AMY), neocortex (NCX), hippocampus (HIP), mediodorsal nucleus of the thalamus (MD), and cerebellar cortex (CBC). This shows higher expression of *TBC1D23* mRNA expression in cerebellum as compared to all other brain regions from the late prenatal period onwards. **c**) Density plot showing the distribution of transcript counts (the number of transcripts produced by a gene as annotated in Ensemble version 72) for all protein-coding genes and *TBC1D23* specifically (red line). **d**) Probability density plots to show the percentage of "Disease" predictions produced by decision tree models for a “test” set of genes enriched for true gene-phenotype associations (dark blue) and a “control” set defined as the set of all protein coding genes (with the exception of those listed in the November 1^st^. 2017 Genomics England release of neurogenetics panels). The “test” set included all genes classified by Genomics England as having moderate (amber) or low (red) evidence for association with a neurogenetic disorder as well as high evidence genes added to a panel after March 31^st^ 2017 and until November 1^st^ 2017. None of the genes used in this “test” set were used for learning the classifier ensembles. There was a clear and highly significant (Mann-Whitney two-sample unpaired test p-value = 2.2×10^−16^) difference in the distribution of “Disease” quality scores comparing “test” and “control” gene sets.

While these findings were encouraging, the limited availability of relevant novel gene-phenotype associations, meant that any measure of prediction accuracy would be unstable. With this in mind, we also investigated gene-phenotype associations considered to have lower levels of supporting evidence, termed “Amber” (borderline) or “Red” (low) genes within the Genomics England PanelApp (PanelApp Handbook Version 5.7). For this list of 329 gene-phenotype associations we calculated the percentage of “Disease” votes (“Disease” quality score) made for each gene by the relevant classifier and demonstrated that these values were highly skewed towards higher values (Figure 6d, **dark blue density plot**). In contrast, the distribution of corresponding values for a control gene set, consisting of all protein coding genes with the exception of those listed in the November 1^st^. 2017 Genomics England release of neurogenetics panels, was negatively skewed (Figure 6d, **light blue density plot)**. The difference in these distributions of “Disease” quality scores was highly significant (Mann-Whitney two-sample unpaired test p-value = 2.2×10^−16^). Given that the list of test genes was likely to be highly enriched for true gene-phenotype associations, this finding provides additional evidence that the classifier ensembles we have generated capture some if not all the key characteristics of disease-associated genes.

## Discussion

The overall aim of this study was to test the hypothesis that genes contributing to Mendelian forms of neurogenetic disorders can be identified using a relatively limited set of gene-based predictors, which do not incorporate directly or indirectly any prior biological knowledge. We achieve this aim through the generation of machine learning classifier ensembles for 25 neurogenetic disorders and their resulting predictions. We demonstrate that our set of disease-specific gene predictions are enriched for GO terms and molecular pathways already implicated in neurogenetic diseases. Furthermore, we show that when mutated in mouse model systems these predicted gene sets produce phenotypes, which are highly enriched for neurological abnormalities and which can mirror the expected human phenotype with accuracy. Finally and most importantly we demonstrate an 8.2 fold enrichment over chance in disease gene predictions for a limited set of recently identified neuro-disease genes. Given that our gene predictions could potentially be used to increase the yield of diagnostic whole exome sequencing by helping investigators prioritise genes of interest, we have made our predicted gene sets available and easily searchable through a web application G2P (Gene 2 Phenotype, accessible at XXXXX).

Moving beyond the predictive value of our ML-based classifier ensembles, we provide evidence of the explanatory power of this approach. In particular, the finding that genes contributing to Mendelian forms of neurogenetic tend to be complex in structure, with a higher than expected number of transcripts and unique exon-exon junctions annotated per gene. Interestingly, this would suggest that these genes could be particularly prone to splicing mutations, which are more difficult to recognise and assess.

Most importantly, and despite the limitations of our approach we demonstrate the value of machine learning approaches in understanding neurogenetic disorders now that a critical mass of genome-wide annotation and gene discovery data exists. Given that both types of data are only set to increase in quality and quantity across a wide range of disorders we would envisage that ML-based approaches of this kind are likely to increase in impact over the next 5 years.

---

**Supplementary Table 1**

Table providing the names and affiliations of all individuals contributing to the curation of Genomics England disease-associated panels used in this study.

**Supplementary Table 2**

Table describing all disease-associated panels used and their contents.

**Supplementary Table 3**

Table providing all predictors used to describe genes with omics.

**Supplementary Table 4**

Table providing r2 and p-values for the relationship between all 154 predictors.

**Supplementary Table 5**

Table showing the predictors which differed significantly between and non-disease genes for each disease-associated panel.

**Supplementary Table 6**

Table providing all disease gene predictions made using ML classifier ensembles with a stringency of 90%.

**Supplementary Table 7**

Table providing significant gene ontology, KEGG and REACTOME pathway enrichments for gene predictions made using disease-specific ML classifier ensembles.

**Supplementary Table 8**

Table providing significant cell type-specific enrichments for gene predictions made using disease-specific ML classifier ensembles.

**Supplementary Table 9**

Table providing significant Mammalian Phenotype Ontology terms for gene predictions made using disease-specific ML classifier ensembles.

**Supplementary Table 10**

Table showing the predictions for 32 new, high confidence gene-phenotype associations for which the relevant ML-based classifier provided a “Disease” or “Non-disease” prediction at a stringency of 0.9.

**Supplementary Figure 1**

**ROC values for all Genomics England panels for each Caret Algorithm used for model learning:** Box and whisker plots to show the distribution of ROC values (x axis) for predictions generated using the training data (each GE panel genes corresponds to a point in the sample) per classifier (y axis). While many algorithms performed well, we chose J48 because it has a good trade off between ROC and explanatory capabilities.

**Supplementary Figure 2**

**ROC values for all Caret Algorithms used for model learning on each Genomics England panel:** In this box and whisker plot we show the distribution of ROC values (x axis) for predictions generated using the training data for a single panel (y axis, each GE panel comes with its size in genes). Each algorithm corresponds to a point in the sample set. The plot shows that the learning ROC performance for all panels show a performance better than random but with room for improvement.

**Supplementary Figure 3**

**Improvement of ROC values through the use of an ensemble of 200 trees versus a single tree:** Box and whisker plots to demonstrate that there is an improvement in ROC values (x axis, % of improvement in ROC) on test data for all GE panels (y axis) when using an ensemble of decision trees as compared to a single tree to identify gene-phenotype associations.

## Online methods

### Identification of genes causing Mendelian neurogenetic disorders

We obtained our list of genes causing Mendelian neurogenetic disorders from the Genomics England Panel App (https://panelapp.genomicsengland.co.uk/; March 31^st^ 2017). We considered all panels included under the level 2 heading “Neurology and Neurodevelopmental disorders”. We further analysed all gene panels with more than 10 “Green” (diagnostic-grade) genes with the exception of “Intellectual Disability” due to the very broad phenotypic spectrum associated with this gene panel. This resulted in the use of 25 gene panels, equating to 1140 unique genes (Supplementary Table 2).

### Identification of control gene sets

Given that we aim to use ML-based classifiers not only to predict new gene-phenotype associations, but to provide explanatory information as well, we require a “control” gene set, which can serve as negative examples that better delineate through contrast the core features of disease genes. We define a control gene, as any gene with no currently known association to any disease (whether neurological in nature or otherwise). In this setting we only consider disease associations based on Mendelian inheritance and accept that some of the genes defined as “control” may be identified in the future as having a disease association. Thus, as in the case of Chakraborty and colleagues^31^, we generate a list of “control” genes by excluding all genes currently contained within the Online Mendelian Inheritance in Man database (https://www.omim.org/), Genetic Association Database (https://geneticassociationdb.nih.gov/) and the Human Gene Mutation Database (http://www.hgmd.cf.ac.uk/ac/index.php) from the Ensembl (version 72) gene set. Using this approach we identify 4260 “control” genes.

### Extraction of gene-based measures of genetic constraint

The Exome Aggregation Consortium (ExAC, http://exac.broadinstitute.org/) database has been used to model the difference between the expected and observed frequency of loss-of-function and missense mutations in all genes. This information has been captured through the gene-specific parameters, ExACpLi (the probability of being loss-of-function, lof, intolerant of both heterozygous and homozygous lof variants), ExACpRec (the probability of being intolerant of homozygous, but not heterozygous lof variants), ExACpNull (the probability of being tolerant of both heterozygous and homozygous lof variants) and ExACpMiss (corrected missense Z score, taking into account that higher Z scores indicate that the transcript is more intolerant of variation). These four measures of genetic constraint were downloaded from the ExAC consortium site (ftp://ftp.broadinstitute.org/pub/ExAC_release/release0.3/functional_gene_constraint/; released January 13^th^ 2017) for use as predictors.

### Extraction and generation of measures of gene complexity

All measures of gene complexity were either extracted directly or generated from Ensembl version 72. The intronic length of each gene (IntronicLength) was determined using an ad-hoc script that calculates intron length in basepairs considering all possible transcripts present in the gene. The total number of unique exon-exon junctions generated by each gene (numJunctions) was calculated using the refGenome R package with all the known junctions summed to obtain a value per gene. The number of genes overlapping a gene of interest (countsOverlap) was estimated using the R package GRanges^32^, such that for each gene the number of genes overlapping 1bp or more with the gene of interest, regardless of strand, was counted. The number of overlapping protein coding genes (countsProtCodOverlap) was generated in a similar manner except that only genes classed as “protein coding” within Ensembl (version 72) were considered.

### Generation of measures of tissue-specific gene expression & co-expression

In order to generate a set of gene-based measures of tissue-specific gene expression and co-expression, we used the GTEx^7^ V6 gene expression dataset (accessible from https://www.gtexportal.org/home/) which comprises 54 tissues and expression data for 8557 samples covering 56318 Ensembl genes. We first filtered the GTEx dataset by tissue sample availability using only those tissues with more than 60 samples available leading to the consideration of 47 tissues. We initially analysed each of the 47 tissue sample sets separately by filtering genes on the basis of an RPKM > 0.1 (observed in > 80% of the samples for a given tissue). We then corrected for batch effects, age, gender and RIN using ComBat^33^. Finally, we used the residuals of these linear regression models to construct gene co-expression networks for each tissue using WGCNA^17^ and post-processing with k-means^18^ to improve gene clustering. Thus, each of the 47 tissues had a corresponding network consisting of a set of gene modules. For each gene in a network we obtained its module membership and adjacency, where Module Membership for a gene g was defined as the correlation of the residual gene expression for gene g and the eigengene of the module it belonged to, and adjacency was defined for a gene g as the sum of all values for its row/column within the Topological Overlap Matrix created by WGCNA.

We used the resulting gene expression and co-expression data to obtain 3 measures of tissue-specific expression. For each tissue any given gene was defined as having tissue-specific expression if the gene expression in that tissue was 3.5 fold higher than the average value across all other 47 GTEx tissues. Similarly, a gene was defined as having tissue-specific module membership or adjacency, if its module membership or adjacency within the co-expression network, was 3.5 fold higher than that across all other remaining GTEx tissues.

### Univariate analysis on single predictors

We wanted to test whether each single predictor had any predictive power on the task of identifying disease genes amongst the whole protein coding genome. For such a purpose, we defined each Genomics England gene panel’s (GP) test dataset as

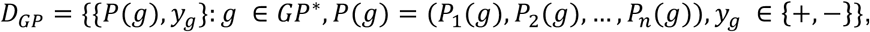

such that GP^*^ refers to a set of genes compound of the genes in the Genomics England GP gene panel plus the rest of the whole protein coding genome (see below), the P_i_(g) is the value of the i-th predictor for gene g and *y*_*g*_ the label of gene *g* for that gene set, {+, −}, where + means gene belonging to the Genomics England panel and – means otherwise. We use predictors of a different nature, namely categorical predictors (including all predictors relating to tissue-specific expression and co-expression) and numeric predictors (including all predictors relating to genetic constraints and gene complexity). Accordingly, we applied two different statistical tests to look for significant differences between disease and non-disease genes. For categorical predictors we used Fisher’s exact tests (FET) with a control gene set consisting of all genes within Ensembl (version 72) with tissue-specific data available and with the exception of genes within the Genomics England “Neurology and neurodevelopmental disorders” panels (17,315 genes). For numeric predictors we used Mann-Whitney U tests (MWU) with a control gene set consisting of all protein coding genes within Ensembl with a non-null value for the predictor and with the exception of genes within the Genomics England “Neurology and neurodevelopmental disorders” panels (17,918 genes). FETs an MWU tests were performed under the null that there was no significant difference in predictor values found in disease gene panels in comparison to the control gene set. We report Bonferroni-Hochberg corrected p-values in all cases.

### Machine leaning to generate disease-specific classifiers

We used a supervised machine learning (ML) approach^34^ to inductively model gene-phenotype associations and so generate classifiers that can distinguish between genes that have an association with a neurological phenotype and those that do not. Formally stated, this meant that given a set of genes already causally linked to a specific neurological phenotype and defined by a Genomics England panel, GP, we sought a classifier C_GP_, such that, for any gene *g*, it predicted whether it too would have a gene-phenotype association and denote it in the form of a function, as follows:

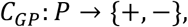

and *P* is a set of *P*_*i*_ predictors such that *P*(*g*) = (*P*_1_(*g*), *P*_2_(*g*), …, *P*_*n*_(*g*)) is a vector of attributes of gene *g* and the output of the classifier is then *C*_*GP*_(*g*) = + if the classifier predicts g to have a gene-phenotype association for phenotype GP and *C*_*GP*_(*g*) = − otherwise.

The global ML analysis is divided into two main phases. In the 1^st^ phase we seek for the best possible learning algorithm we have available given requirements on prediction accuracy (mainly ROC values on 10-fold cross-validation) and model interpretability. In the 2^nd^ phase, we use a high number of such models into a voting based integration ensemble to account for the acute imbalance in the proportion of positive and negative examples in the learning data.

1^st^ phase: identification of the best ML algorithm

Using the Caret R package^19^ we constructed simple classifiers *C*_*GP*_ for 14 ML techniques including the main supervised learning paradigms and all the 25 Genomics England working panels. The learning data for each panel was compound by all the High Evidence disease genes belonging to the panel and all the Chakraborty control genes, 4260. The Caret algorithms used were: (1) trees/rules based algorithms (C5.0Tree, LMT, J48, ONeR, Rpart), (2) neural network based algorithms (NNet, GlmNet), (3) linear regression based algorithms (PLR, RRLDA, svmLinear), a bayes reasoning approach with NaiveBayes and (5) ensemble based algorithms (AdaBoost, Boruta, Rborist). We evaluated each Caret candidate ML algorithm on each *C*_*GP*_ dataset using repeated cross-validation (10 fold) and automatic hyperparameter evaluation as provided by Caret. ROC, specificity and sensitivity for all Caret algorithms applied to all Genomics England panels were evaluated (Supplementary Figure 1, results segregated by algorithm; Supplementary Figure 2, results segregated by Genomics England panel). While Rborist and Boruta showed significantly better ROC values than J48 (P < 0.0008 and P < 0.02 respectively with a paired t-test), J48 still had a high predictive power (only 9-10% lower than Rborist and Boruta). Given that this algorithm had much greater explanatory power, all subsequent analyses were performed with J48.

2^nd^ phase: construction of the best predictor based on J48

We used the J48 algorithm within a wrapper (i.e. an ensemble in ML terminology) that we developed in order to deal with the disproportionate number of negative examples in our learning set. This wrapper involved the generation and integration of many ML models from multiple learning data sets, all with the same set of positive examples, but with different negative examples generated through random re-sampling of the Chakraborty control gene set. Formally stated, this meant that given a panel *GP*, and its learning data defined as:

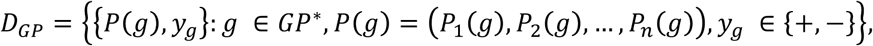

such that GP^*^ refers now to a set of genes compound of the genes in the Genomics England GP gene panel as positive examples plus the Chakraborty genes as negative ones, we created (*D*_*GP*1_, *D*_*GP*2_, …, *D*_*GP*200_) learning datasets, with identical predictors *P* but now, the {*y*_*g*_} had a 1:1 ratio of positive (+) and negative examples (−). The set of positive examples in each learning dataset was identical to that in *D*_*GP*_, but the negative examples were samples with replacement from the Chakraborty control genes.

We applied J48 to each of the learning datasets (*D*_*GP*1_, *D*_*GP*2_, …, *D*_*GP*200_) to obtain (*C*_*GP*1_, *C*_*GP*2_, …, *C*_*GP*200_) classifiers, which we aggregated into a single classifier, *C*_*GP*_ = Δ(*C*_*GP*1_, *C*_*GP*2_, …, *C*_*GP*200_), where Δ was a function to integrate responses from all individual classifiers. Δ has to be designed taking into account that the ensemble compound of the 200 ML models with have a limited predictive power for a number of reasons. Firstly, the number of disease genes to learn from will be limited, and most likely incomplete, for all panels. And secondly, likely amongst the genes we consider as controls there will be disease genes we do not know yet. Therefore, we must come up with a strategy that deals with that. And we define a stringency parameter to control the eagerness with which the ensemble generates a prediction, i.e. Disease or Non-disease but also to include a third outcome: Uncertain. And we define a parameter called stringency, let it be denoted with s in [0,1] that works in the following way: given a gene g, and the 200 models generated from a gene panel GP, we generate 200 predictions *C*_*GPi*_(*g*), *i* = 1, …, 200. Either Disease or Nondisease. Let p_d_ and p_n_ the proportion of Disease and Non-disease predictions for g, such that p_d_ + p_n_ = 1. If p_d_ > p_n_, the outcome of Δ(*C*_*GPi*_(*g*), *i* = 1, …, 200) will be Disease when p_d_ > s. Analogously when p_n_ > p_d_. The outcome will be Uncertain otherwise.

### Testing for annotation term enrichment amongst predicted genes

We used gProfileR^35^ to investigate enrichment of Gene Ontology, REACTOME and KEGG pathways annotation terms amongst predicted gene sets. We included IEA (Inferred Electronic Annotations) and used the gSCS test developed by the authors to assess for annotation term enrichment. The graphical representation of the REACTOME term enrichment was based on the most significant terms as reported by gProfileR and by using Cytoscape 3.5^36^. The enrichment of Mammalian Phenotype Ontology terms amongst mouse models with mutations in mouse orthologues of predicted disease genes was performed using the ToppFun function within ToppGene (https://toppgene.cchmc.org/, REF) and applying the default settings.

### Testing for enrichment of genes expressed in a cell-specific manner amongst predicted genes

We obtained cell-type specific gene lists relevant to brain from Soreq et al.^26^. These lists were based on the analysis of RNA-sequencing data from purified cells isolated from mouse cerebral cortex ^37^and covered astrocytes, neurons, oligodendrocyte precursors, newly formed oligodendrocytes, myelinating oligodendrocytes, microglia and endothelial cells. Genes that appeared in more than one cell type-specific gene list were removed from the analysis, and the remaining genes were converted to human orthologs using the biomaRt package^38^. We used the Fisher’s Exact Test to check for enrichment of cell-specific markers with each set of predicted genes. The false discovery rates (FDR) were calculated using the Benjamini-Hochberg procedure^39^ and an FDR threshold of 5% was used to assess the significance of the enrichment observed.

### Assessing the importance of predictors in disease-specific classifier ensembles

Given a set of trees from an ensemble, we measure the importance of each ML predictor within the ensemble on the basis of its relevance when classifying genes as “Disease”. We base the calculation of importance on two different properties of a predictor when used within a set of trees, namely the frequency of appearance across the trees and the average depth of the predictor within each tree. Clearly, when a predictor appears high in a tree (i.e. near to the root or even the root itself) and its subtree leads to a higher proportion of genes classified as “Disease”, the predictor has relevance for disease classification. Moreover, when the attribute is used many times across trees it is also important. We calculate two measures to reflect these properties. Note that each non-leaf node on a tree is a test (positive test in this case) for a predictor. Leaf nodes are labels, either “Disease” or “Non-disease”. In regard to depth, given a predictor p and tree t, with a maximum of n nodes between the root and any leave, *depth* is defined for the predictor p in the tree t as the mean of the p_1_, p_2_, …, p_m_ values when p appears m times at the tree t and p_i_ is n minus the number of nodes from the root to that node at the i-th appearance of the predictor within the tree. Depth is always in [0,1]. With respect to appearance, the appearance of a predictor p in a trees ensemble is simply its frequency of appearance as root node of a subtree with higher proportion of genes classified as “Disease” for that subtree generating a value in [0,1]. As a single measure, we simply add these two measures. As a single measure we simply add these two values.

### Code availability statement

The code we have developed for this paper is available right now upon request.

We are working on making it easier to use. It will be ready before publication.

### Data availability statement

All the new data produced in this paper has been made available through the supplementary materials accessible from the submission.

#### Acknowledgements

Mina Ryten, David Zhang and Karishma D’Sa were supported by the UK Medical Research Council (MRC) through the award of Tenure-track Clinician Scientist Fellowship to Mina Ryten (MR/N008324/1). Sebastian Guelfi was supported by Alzheimer’s Research UK through the award of a PhD Fellowship (ARUK-PhD2014-16). Regina Reynolds was supported through the award of a Leonard Wolfson Doctoral Training Fellowship in Neurodegeneration.

Henry Houlden was supported by the UK Medical Research Council, the Wellcome Trust, Muscular Dystrophy UK, Muscular Dystrophy Association USA and the National Institute for Health Research.

This research was made possible through access to the data and findings generated by the 100,000 Genomes Project. The 100,000 Genomes Project is managed by Genomics England Limited (a wholly owned company of the Department of Health). The 100,000 Genomes Project is funded by the National Institute for Health Research and NHS England. The Wellcome Trust, Cancer Research UK and the Medical Research Council have also funded research infrastructure. The 100,000 Genomes Project uses data provided by patients and collected by the National Health Service as part of their care and support.

## Author contributions

Juan Botía generated the machine learning classifier ensembles. Juan Botía, Sebastian Guelfi, David Zhang, Karishma D’Sa, Regina Reynolds and Mina Ryten assessed and analysed the output of classifier ensembles. Daniel Onah and Juan Botía designed and implemented a web server to access predictions.

Ellen McDonagh and Antonio Rueda Martin conceived, designed, built and created PanelApp where 800 registered reviewers worldwide have supplied data or curated 167 panels for genome analysis and helped establish 213 reviewable panels with 4104 genes. Augusto Rendon contributed to the design of PanelApp and coordinated the development and curation efforts.

Arianna Tucci and Henry Houlden (on behalf of the Neurology Genomics England Clinical Interpretation Partnership) assisted and led the curation of gene panels.

Juan Botía, David Zhang, John Hardy and Mina Ryten prepared the manuscript. Juan Botía and Mina Ryten conceived and designed the project.

